# Fibrillin microfibril structure identifies long-range effects of inherited pathogenic mutations affecting a key regulatory TGFβ-binding site

**DOI:** 10.1101/2022.08.26.505362

**Authors:** Alan RF Godwin, Jennifer Thomson, David F Holmes, Christin S Adamo, Gerhard Sengle, Michael J Sherratt, Alan M Roseman, Clair Baldock

**Affiliations:** Wellcome Centre for Cell-Matrix Research, Division of Cell-Matrix Biology and Regenerative Medicine, School of Biological Sciences, Faculty of Biology, Medicine and Health, University of Manchester, Manchester Academic Health Science Centre, Manchester, M13 9PT, UK; Center for Biochemistry, Faculty of Medicine and University Hospital Cologne, University of Cologne, Cologne, Germany; Department of Pediatrics and Adolescent Medicine, Faculty of Medicine and University Hospital Cologne, University of Cologne, Cologne, Germany; Center for Molecular Medicine Cologne (CMMC), University of Cologne, Cologne, Germany; Cologne Center for Musculoskeletal Biomechanics (CCMB), Cologne, Germany; Division of Cell-Matrix Biology and Regenerative Medicine, School of Biological Sciences, Faculty of Biology, Medicine and Health, University of Manchester, Manchester Academic Health Science Centre, Manchester, M13 9PT, UK; Division of Molecular and Cellular Function, School of Biological Sciences, Faculty of Biology, Medicine and Health, University of Manchester, Manchester Academic Health Science Centre, Manchester, M13 9PT, UK

## Abstract

Genetic mutations in fibrillin microfibrils cause a range of serious inherited diseases such as Marfan syndrome (MFS) and Weill-Marchesani syndrome (WMS). These diseases typically show major dysregulation of tissue development and growth, particularly in skeletal long bones, but links between the mutations and the diseases are unknown. In this study we reveal the detailed cryo-EM structure of native fibrillin microfibrils from mammalian tissue. The major bead region showed pseudo 8-fold symmetry and a buried protease resistant N-terminal core. Based on this structure, we show a WMS deletion mutant induces a rearrangement with long-range effects blocking interaction with latent TGFβ-binding protein (LTBP)-1 at a remote site. Separate deletion of this binding site resulted in the assembly of shorter fibrillin microfibrils with structural alterations. These results establish that in complex extracellular protein assemblies, such as in fibrillin, mutations may have long-range structural consequences to disrupt growth factor signalling and cause disease.

## Introduction

Elastic fibres are essential components of all mammalian elastic tissues such as blood vessels, lung, joints and skin [1]. The main components of elastic fibres are elastin and fibrillin, whereby elastic fibre formation requires fibrillin microfibril as a scaffold for the correct deposition of tropoelastin. Fibrillin microfibrils also provide limited elasticity in tissues devoid of elastin such as the ciliary zonule, an elastic ligament essential for lens attachment in the eye. In many tissues, these assemblies provide a multifunctional platform for the interaction of matrix molecules required for elastic fibre assembly and function, and provide a connection to the cell surface [2]. Fibrillin is needed for the correct assembly of many microfibril associated proteins, including members of the latent transforming growth factor (TGF)-β binding protein (LTBP) family, and also mediates interactions between microfibril binding proteins facilitating their functions [2].

The fibrillin superfamily is composed of fibrillins (isoforms 1-3), with fibrillin-1 being the predominant form [3–6], and the four structurally related LTBPs (isoforms 1-4) [7–10]. Fibrillin superfamily proteins are composed primarily of arrays of epidermal growth factor-like (EGF) domains, interspersed with TGFβ-binding like (TB) domains and hybrid domains [3]. The three fibrillin isoforms are highly homologous to each other with differences including a proline rich region in fibrillin-1, which in fibrillin-2 is glycine rich, and in fibrillin-3 is proline and glycine rich. Of the 47 EGF domains in fibrillin, 43 are calcium binding (cb) [3]. There are seven TB domains (also referred to as 8-cysteine domains) which are unique to the fibrillin superfamily.

Fibrillin assembles to form beaded microfibrils with a ~56 nm periodicity [11] and a mass of ~2.55 MDa per repeat [12]. These microfibrils are polar polymers which are formed by linear assembly of fibrillin molecules via direct interactions between the N- and C-termini [13]. Lateral association of fibrillin molecules also occurs and is driven by a homotypic interaction between these termini to form mature microfibrils [14–16]. Microfibril structure can be described by three distinct regions based on their banding pattern, these have been termed the bead, arms and interbead regions (see Figure 1B) [17, 18]. However, how fibrillin molecules are organised into mature microfibrils and their 3D structure are still unclear, as their complexity and variability prevent their analysis by most structural biology and biochemical tools. Therefore, a number of packing models have been proposed, including an intramolecular pleating model, where each fibrillin molecule spans one 56 nm period [19] and a one-half staggered model, where each fibrillin molecule spans two 56 nm periods [20]. Both models suggest that the N- and C-termini are located near the bead region.

**Figure 1.**
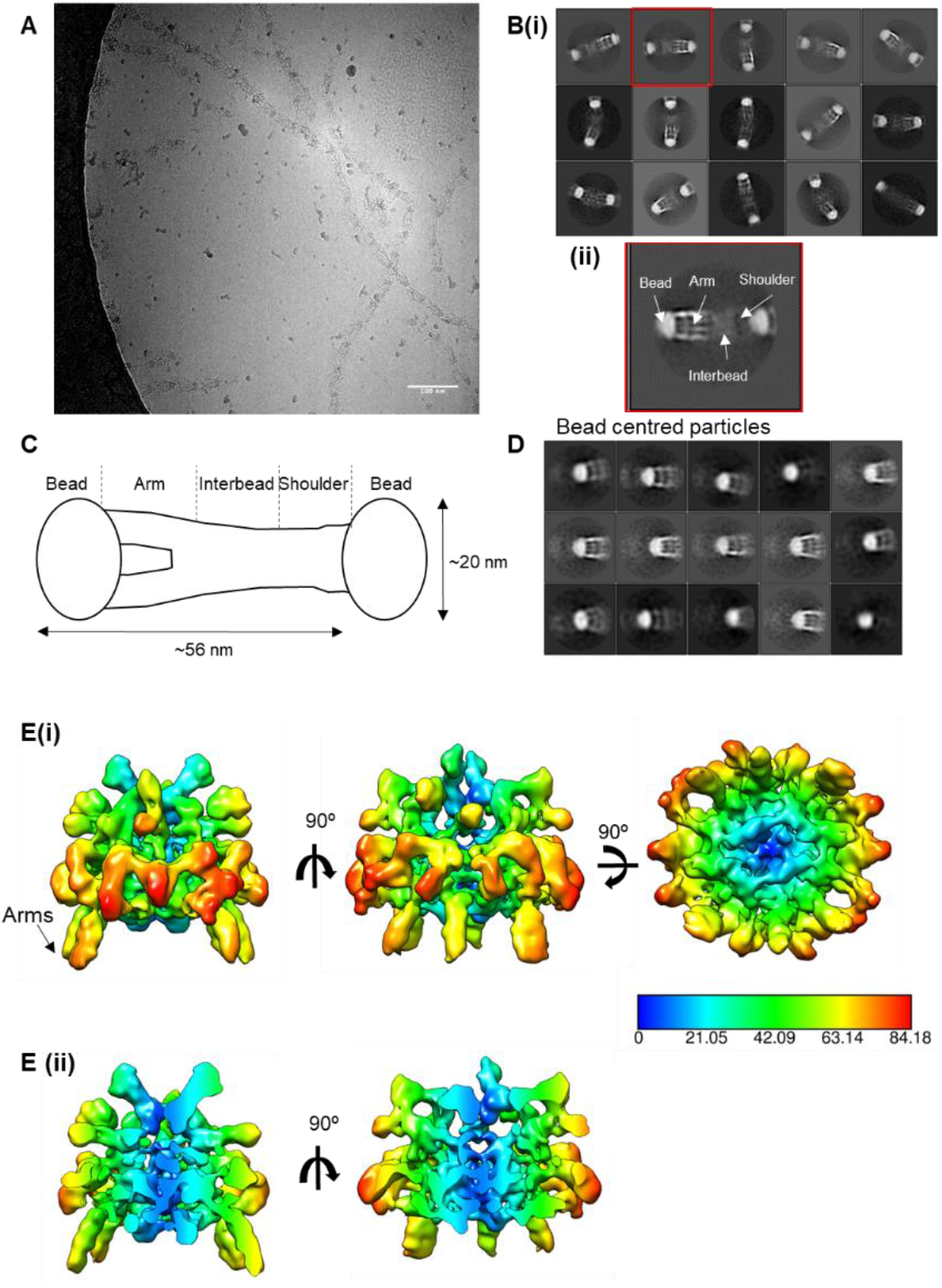
Cryo-EM structure of the Fibrillin microfibril bead region. Fibrillin microfibrils were extracted from bovine ciliary zonules in non-denaturing conditions and imaged using cryoEM. A) Representative cryoEM image of purified bovine ciliary zonule microfibrils. Scale bar = 100 nm. B(i) Reference free classification of the full fibrillin microfibril repeat. Box size = 100 nm. B(ii) A class highlighted in panel (i) with a red box, was rotated 180° and enlarged to highlight the different microfibril regions. C) A cartoon schematic of the fibrillin microfibril highlighting the bead, arm, interbead and shoulder regions of the microfibril. D) Classification of particles aligned to the fibrillin bead region. Box size = 57 nm. E) The cryoEM single particle reconstruction of the fibrillin bead region shown in three orthogonal orientations. The bead is rainbow coloured by cylindrical radius from blue, at the centre of the bead, to red on the outside of the bead. The colour key is shown with radius in Å. ii) Two orthogonal views of the bead reconstruction have been sliced to show a cross-section through the centre of the bead.

In addition to their role in elastic fibre assembly, fibrillin microfibrils have a key role in tissue homeostasis, which when perturbed by fibrillin mutation causes a number of heritable connective tissue disorders such as Marfan syndrome (MFS) and Weill Marchesani syndrome (WMS) [21]. The role of fibrillin in tissue homeostasis is mediated via its interactions with cell surface receptors, such as integrins [22, 23] and syndecans [24], and with growth factors such as TGFβ and bone morphogenetic proteins (BMPs) [25–27]. Of the four LTBP isoforms, LTBP-1, −3 and −4 play important roles in the processing and secretion of TGFβ [28], through covalently binding to the latency-associated peptide of TGFβ (LAP), producing the large latent TGFβ complex (LLC) [29, 30]. The LLC becomes sequestered within the matrix, and can regulate TGFβ bioavailability [29–31]. Fibrillin is thought to have a role in the regulation of latent TGFβ as the first hybrid domain in fibrillin binds to the C-terminal region of LTBP-1 [25] and dysregulated TGFβ signalling is a characteristic of MFS [32]. However, mice homozygous for the deletion of the fibrillin-1 hybrid domain are able to form normal microfibrils [33]. A common WMS mutation is a three-domain fibrillin-1 deletion in an adjacent region (domains TB1-proline-rich region-EGF4). This deletion when replicated in mice resulted in a WMS-like phenotype with thick skin, short stature, and brachydactyly [34].

Although growth factor binding and disease-linked sites have been mapped *in vitro* using recombinant fibrillin-1 fragments, their exact sites of interaction on the microfibril are still unknown. Most microfibril binding proteins including proBMP complexes, LTBPs (and the LLC), fibulins, microfibril associated glycoproteins (MAGPs), and ADAMTS/L proteins bind near to the N-terminal region of the fibrillin monomer (for review see [2]). However, it remains to be elucidated how multiple binding partners can interact with microfibrils and whether there is competition for these interactions in the assembled microfibril. Previous negative stain electron microscopy studies have not achieved sufficient resolution (~43 Å) to determine the arrangement of fibrillin molecules in tissue microfibrils [35]. To address this lack of structural information, we have taken advantage of the recent advances in cryo-electron microscopy to determine the structure of fibrillin microfibrils extracted from mammalian tissue to sub-nm resolution. This structure allows for the first time the tracking of individual fibrillin molecules through the bead and interbead regions. To confirm the location of important functional regions within the microfibril, microfibrils from mouse models containing deletions of the first hybrid domain that binds latent TGFβ, and the WMS-causing deletion were analysed. The microfibrils have reduced periodicities and structural perturbations, which together with binding analyses and STEM mass mapping have defined the fibrillin molecular organisation, and the location of the LTBP-1 (and LLC) binding site.

## Results

### 3D reconstruction of the fibrillin microfibril bead region

To determine the 3D structure of the fibrillin microfibril repeat, microfibrils were purified from ciliary zonules using sonication and size exclusion chromatography and imaged using cryoEM (Figure 1A). When imaged, in a frozen native hydrated state, the microfibrils had the characteristic beads-on-a-string appearance [35, 36]. 27,737 microfibril repeating units were digitally extracted from the images and classified into 2D classes (Figure 1B). Some heterogeneity is apparent in the class averages due to flexibility in the microfibrils along the microfibril axis, as we have described previously [35]. Therefore, to overcome this and with the objective of increasing the resolution of the microfibril reconstruction, separate sub-models of specific microfibril regions (i.e., the bead and arm regions) were created, by centring the reconstruction locally on each of these regions. Particles which had been aligned to the full fibrillin repeat were recentred to locate the bead coordinates at the particle centre using a custom python script. Classification centred on the bead region provided classes that showed the characteristic bead with arms emerging from one side of the bead (Figures 1C and D). We saw four defined regions in 2D class averages of the repeat unit in the cryoEM images termed the bead, arms, interbead and shoulder regions (Figure 1B(ii)). Two-fold symmetry was applied along the microfibril axis and a 3D reconstruction of the bead region was generated in RELION with 7,139 particles used in the final refinement. The resolution of the microfibril bead region was estimated using Fourier shell correlation (FSC) of two independently refined half maps using the 0.143 criterion to give a resolution of 9.7 Angstrom (Supplementary Figure 1). The bead shape is approximately spherical; it can be enclosed by an ellipsoid with dimensions of 16.5 x 15 x 12 nm and has a complex interwoven arrangement (Figure 1E). A mask around the bead was used in the processing but despite not being under the mask, the arm region can still be seen protruding from the bead. The resolution allows the individual fibrillin molecules to be visualised interwoven through the bead. The tight packing within this region supports data that show resistance of the microfibril bead region to proteolysis [37].

### Identifying protease-resistant domains within the bead region

In order to identify the fibrillin domains within the bead region, previous antibody mapping and MS studies were revisited. The binding sites of two monoclonal antibodies, raised against the N-terminal half of the fibrillin molecule, have previously been mapped on the fibrillin microfibril structure [13, 36]. Mab 1919 binds within the arm region, whereas mAb 2502 binds on the shoulder side of the bead. Using short recombinant fibrillin constructs [19], we narrowed down the binding sites of these antibodies by western blotting (Figure 2A and B). The epitope for mAb 2502 was refined considerably as it detected two overlapping fibrillin constructs which restricted the epitope to within domains TB1 and the proline-rich region (PRR). Whereas, mAb 1919 detected a construct containing domains cbEGF7-Hybrid2, immediately downstream of TB2 (Figure 2A). Taken together, these data suggest that TB1 and/or the PRR are located on the peripheral edge of the bead, whereas domains downstream TB2 are found in the interbead region. Our recent MS data identified a protease-resistant fibrillin region comprised of domains EGF4-TB2 [38]. Peptides from this region were never detected by LC-MS/MS after elastase digestion of microfibrils from either skin, fibroblast cultures or ciliary body (Figure 2C). These findings are consistent with an earlier proteomics study on microfibrils from ciliary zonules [39]. Taking this into account together with the antibody epitope mapping data, we suggest that the EGF4-TB2 domains are buried in the bead core and thus protected from proteolytic digestion (Figure 2D).

**Figure 2.**
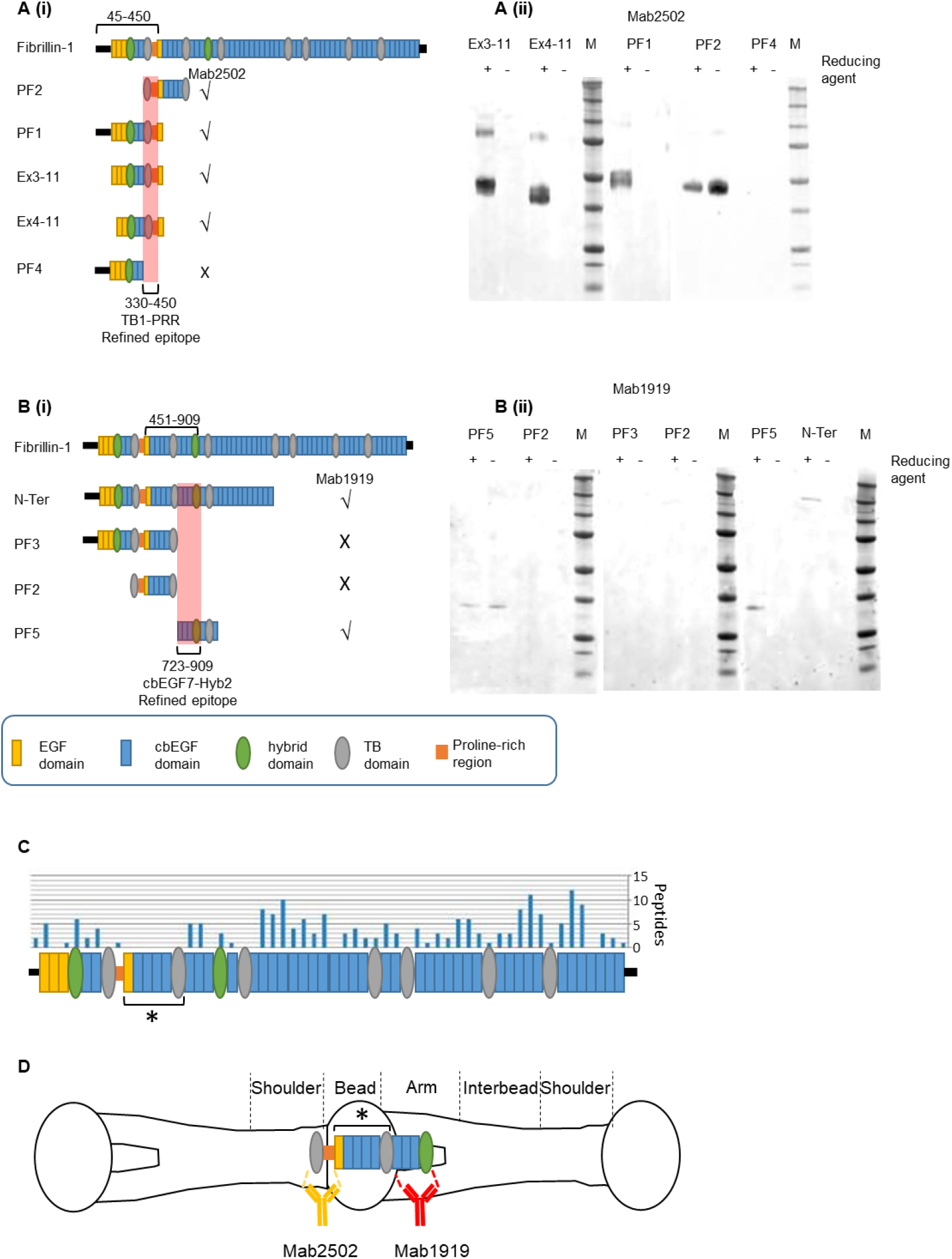
Refining epitope labelling of fibrillin recombinant fragments. Mab25O2 (clone 26 [13]) and Mab1919 (clone 11C1.3) were used to probe overlapping recombinant fibrillin fragments to more accurately define their epitopes in fibrillin-1. (Ai and Aii) schematic diagrams of recombinant fragments N-Ter, PF1, PF2, PF3, PF4, PF5, Ex3-11 and Ex4-11. Recombinant fibrillin-1 fragments after separation by SDS-PAGE in the presence or absence (+/-) of a reducing agent followed by western blotting with either (Aii) Mab2502 or (Bii) Mab1919. The antibody epitopes for mab2502 (45-450 [13]) and mab1919 (451-909) are narrowed down to TB1-PRR (residues 330-450) and cbEGF7-Hyb2 (residues 723-909) respectively, highlighted in red. (C) Number of peptides identified by LC-MS/MS from each domain of fibrillin, where a protease resistant region is located between TB1 to TB2 (TB domains are numbered) and indicated by an asterisk [38]. (D) Diagram of the fibrillin microfibril repeating unit with the binding sites of Mab2502 and Mab1919 on the microfibril (as determined in [36]) and putative location of the protease resistant region identified in [38].

## Modelling the fibrillin N-terminal region into the fibrillin bead region

To try to identify this protease resistant region within the bead, the cryo-electron microscopy (cryoEM) reconstruction was analysed for regions that were buried in the core of the interwoven bead structure. A feature was observed in the centre of the bead map that remained connected at high threshold levels (Figure 3A). This feature can be traced through the centre of the bead and has a persistence length of 17 nm (measured in UCSF Chimera using volume tracer), where the diameter of ~2 nm is consistent with the width of EGF and TB domains and the length supports a chain of ~7 domains. Within the bead, there are four non-symmetric copies of this chain shown in Figure 3B, these four copies appear twice due to the 2-fold symmetry, confirming a pseudo 8-fold symmetric arrangement of eight fibrillin monomers per repeat within the fibrillin microfibril [35, 40]. Three of these features are of similar dimensions but the fourth is shorter. This may be an artefact of the image analysis where some density is lost in the averaging due to the flexibility of the arms. Moreover, the arm region was not under the 3D mask during refinement of the bead, perhaps resulting in the loss of this density. The interface between the bead and arm can be seen more clearly in a lower resolution reconstruction where a larger mask is used for refinement (Supplementary Figure 2). Guided by the antibody-mapping data and starting at the edge of the bead, domain TB1 followed by the consecutive domains the PRR to cbEGF6, were docked into the density for one of these regions. A SAXS derived model of this region was used to aid this docking (Figure 3C), where the arrangement of these domains in the cryoEM structure is very similar to the model based on experimental SAXS data [41].

**Figure 3.**
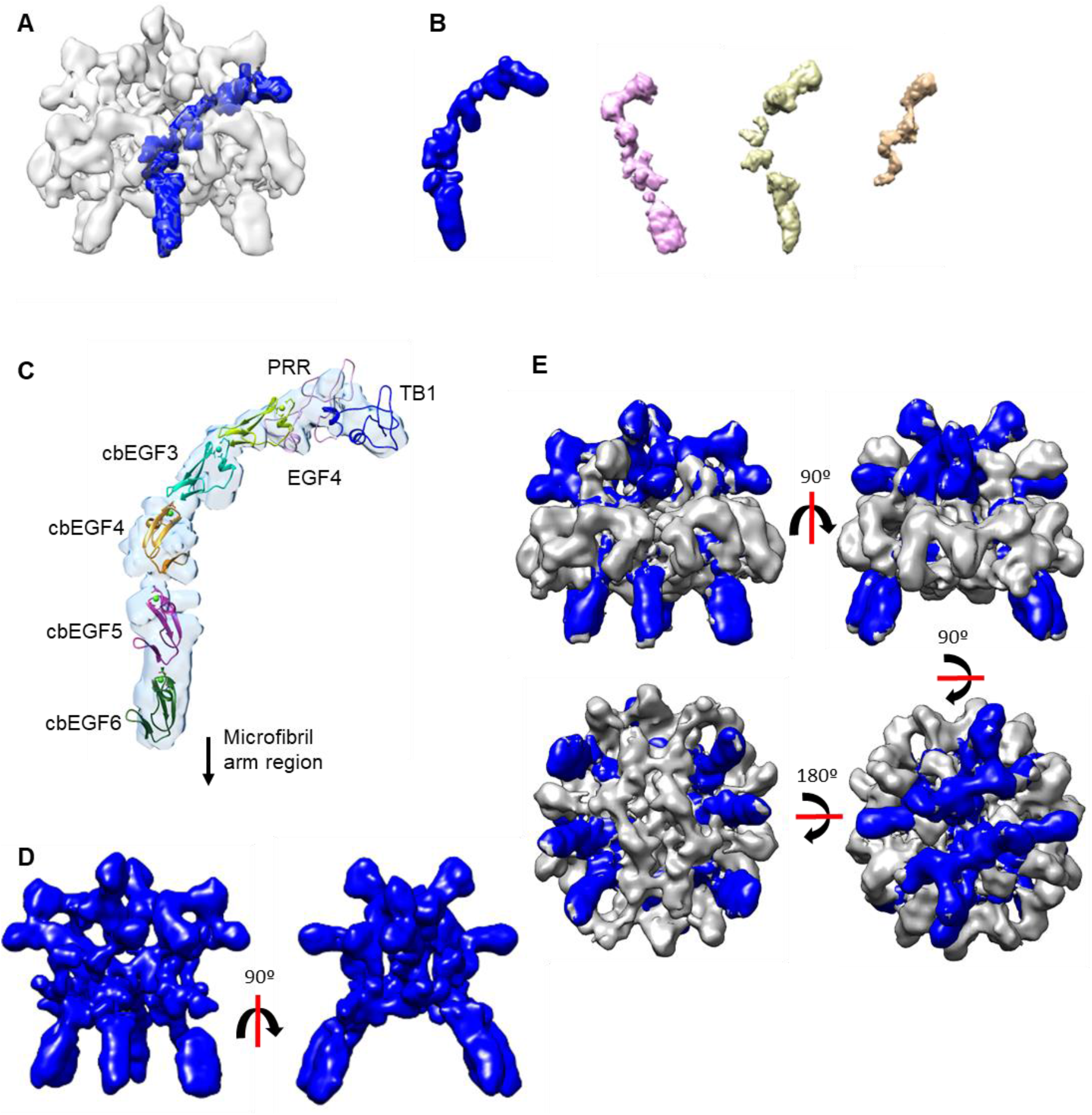
Modelling the fibrillin N-terminal region into the fibrillin bead region. (A) The cryoEM structure of the fibrillin bead region with a feature, in the centre of the bead map, that remained connected at high threshold levels, segmented and highlighted in blue. (B) The four unique (not symmetry related) copies of this region are segmented from the bead map, the feature shown in (A) is coloured blue. As 2-fold symmetry has been applied to the reconstruction, there are also symmetric pairs of each of these features to give eight in total in the bead core. (C) A SAXS derived model of the region encompassing TB1, the PRR, EGF4 and cbEGF3-6 [41] is docked into the region from the core of the bead segmented in (A) (shown as pale blue density). (D) The four N-terminal regions shown in (A) and their symmetric pairs are segmented from the bead structure and coloured blue and are shown without the remaining bead structure. (E) The eight copies of the N-terminal region segmented in blue as shown in (D) with additional bead density shown in grey.

After accounting for the eight copies of this protease-resistant region within the core of the microfibril bead (Figure 3D), the remaining bead structure forms a ring around the outside surface of the bead, as shown in grey in Figure 3E. Antibody binding data have previously shown that the C-terminal region of fibrillin is also located near to the bead in addition to the N-terminal domains [13, 36]. Therefore, this density could correspond to the C- and/or N-termini, which is consistent with microfibril packing models.

## Mutant fibrillin microfibrils reveal disrupted bead region

After locating the domains downstream of TB1 in the bead region, we predicted the upstream N-terminal domains would be in the shoulder region. To confirm this, microfibrils from two fibrillin mouse models with either a domain deletion upstream, or downstream including the TB1 domain, were analysed. Imaging these microfibrils would allow us to confirm the positions of these domains in the microfibril reconstruction and to analyse the structural consequences of deletion of fibrillin domains on microfibril structure. Microfibrils were purified from skin from 6 week old mice homozygous for either a deletion of the first hybrid domain (ΔH1) [33], a domain that contributes to the fibrillin binding site for latent TGFβ via LTBP-1 [25], or a deletion of three domains (TB1-PRR-EGF4) that causes WMS [34]. As microfibrils are more difficult to extract from skin, negative staining EM was used to image microfibrils, as it has a much higher signal-to-noise ratio than cryoEM and can gain more information from fewer microfibrils. 100 microfibril periods were measured for each dataset and compared to wildtype microfibrils (Figure 4A and B). The control microfibrils had a periodicity of 57.95 ± 0.22 nm. The ΔH1 microfibrils had a periodicity of 57.14 ± 0.27 nm, which was 0.81 nm shorter than that of the control. Whereas, the WMS microfibrils had a periodicity of 55.21 ± 0.26 nm which was 2.75 nm shorter than that of the control. These data show that although microfibrils are formed with their main features preserved, the deletion of either one or three domains reduces microfibril periodicity proportionally.

**Figure 4.**
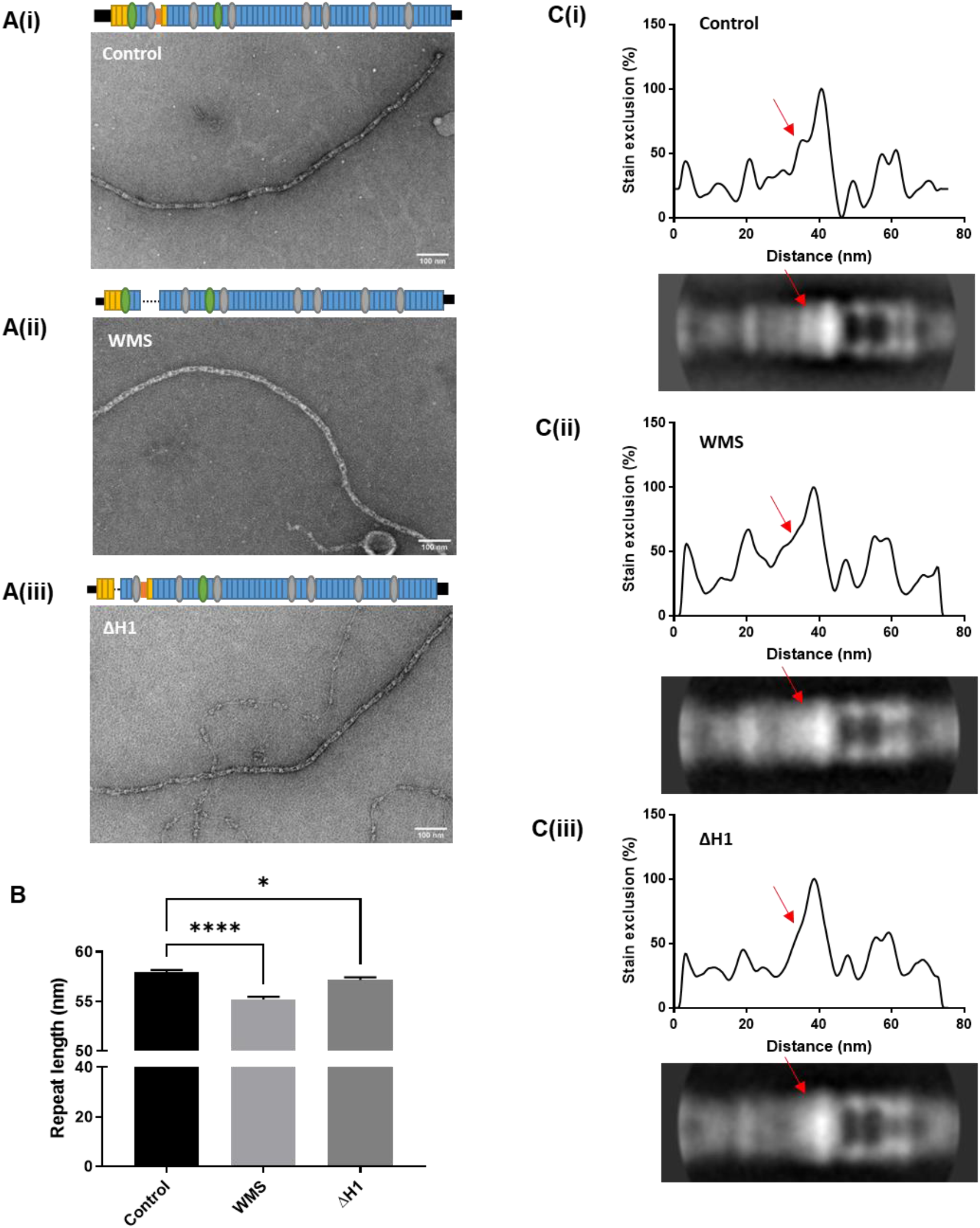
Mutant fibrillin microfibrils reveal disrupted bead region. Fibrillin microfibrils extracted from control and homozygous WMS and ΔH1 mouse skin and imaged using negative stain TEM. (A) Representative TEM images of i) control ii) WMS and iii) ΔH1 extracted fibrillin microfibrils. (B) Bar chart showing the mean periodicity of the control, WMS and ΔH1 extracted fibrillin microfibrils. Error bars are standard error of the mean (SEM). For each data set 100 microfibril periods were measured and statistical analysis shows that the mutant microfibrils are significantly different to the control (* indicates P < 0.05; ****P < 0.0001 as determined by 1-way ANOVA). (C) Averaged images were generated from 458 wildtype periods, 472 ΔH1 periods and 476 WMS periods. The top panel shows a 2D plot of the stain exclusion/intensity profile across averaged fibrillin microfibril repeat period (bottom panel) for (i) control (ii) WMS and (iii) ΔH1 microfibrils. The red arrows highlight a region at one side of the bead of the microfibril which is disrupted in the mutant microfibrils.

Averages from 458 wildtype periods, 472 ΔH1 periods and 476 WMS periods of the microfibril repeating units were made and analysed for changes in microfibril structure. In both cases for the mutants, structural features in the shoulder region are lost (Figure 4C), with the shoulder to the bead peak (seen in the control at ~35 nm) being no longer distinct in WMS and ΔH1 microfibrils. This feature is immediately adjacent to the modelled location of the fibrillin domains through the bead, as shown in Figure 3C, and the location of the mAB2502 epitope [36]. These findings support the location of the TB1 domain in this region, and the docking of these domains through the bead region. We noted that for both deletions, the perturbation spanned a few nanometres suggesting that the deletions may structurally perturb the conformation of neighbouring domains.

## LTBP-1 binds to the bead region of fibrillin microfibrils and its binding is perturbed by a WMS causing mutation

Given the long-range conformational disruption observed in the mutant microfibrils, we predicted that the WMS deletion of domains TB1-PRR-EGF4 could perturb the interaction of microfibril binding proteins that bind near to this deletion. To test this, we analysed the binding of LTBP-1 which interacts with the N-terminal region of fibrillin via the hybrid1 domain [25]. Therefore, to test whether the structural rearrangements, observed in the WMS microfibrils, could impact on the interactions of fibrillin-binding partners, we analysed the binding of the C-terminal region of LTBP-1 to a fibrillin construct containing the WMS deletion. Surface plasmon resonance binding data show that the affinity of LTBP-1 for fibrillin is reduced with the WMS-deletion (Figure 5A and B). However, as these domains are known to not directly be involved in the interaction with LTBP-1 [25], this further supports longer-range structural rearrangements occurring when these domains are deleted.

**Figure 5.**
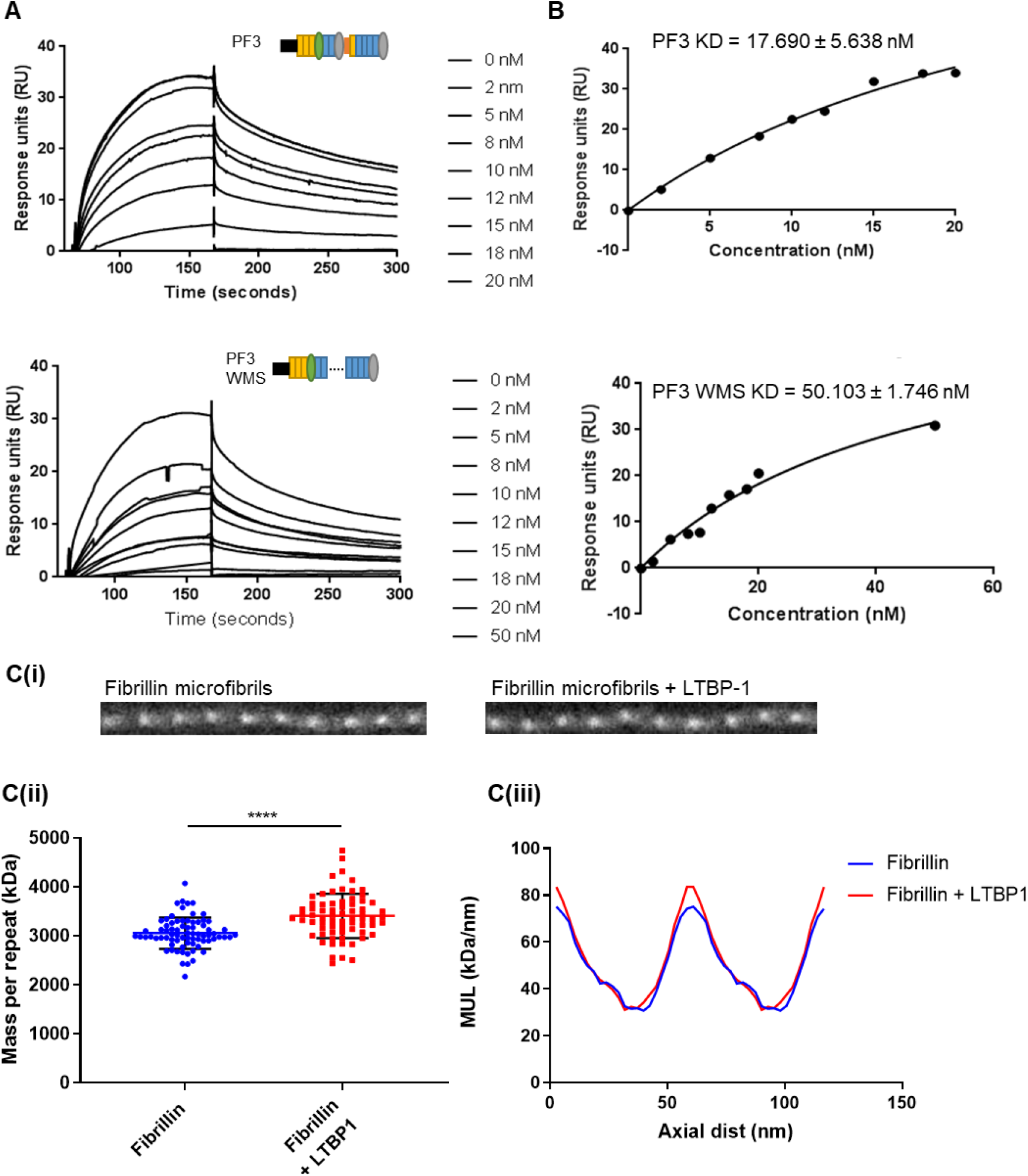
LTBP-1 binds to the bead region of fibrillin microfibrils and is disrupted by a WMS causing mutation. (A) SPR analysis of LTBP-1 binding to the N-terminal fragment PF3 of fibrillin with and without a WMS causing mutation. The LTBP-1 C-terminal region was immobilised to the sensor chip using amine coupling, and concentrations of PF3 (0–20 nM) and PF3 WMS (0–50 nM) were flowed over as analytes. Representative sensorgrams showing binding of PF3 or PF3 WMS to LTBP-1. This experiment was repeated three times. (B) The binding kinetics were determined using equilibrium analysis for the interaction of PF3 or PF3 WMS with LTBP-1. (C) A complex of full length LTBP-1 with purified fibrillin microfibrils was formed and analysed using STEM mass mapping. (i) STEM images of fibrillin microfibrils with and without LTBP-1. (ii) A scatter plot of the mass per microfibril repeat of fibrillin microfibrils with and without LTBP-1. The mean mass of a fibrillin repeat was 3055 ± 36.7 kDa, n=75 from 10 images, in complex with LTBP-1 the mass was 3405 ± 54.43 kDa, n=69 from 11 images. A t-test of the two data sets shows that they are significantly different with a p value of <0.0001. (iii) A trace of the mass per unit length (MUL) across the fibrillin repeat shows that there is a gain in mass at the bead region of the microfibril in the presence of LTBP-1. Each trace is an average of 50 periods (10 measurements from each of 5 images).

Furthermore, given the domain mapping (Figure 3C) and analysis of the ΔH1 microfibril structure (Figure 4C), we hypothesised that LTBP-1 would bind close to the bead region of the microfibril. Therefore, we bound full-length LTBP-1 to purified fibrillin microfibrils and looked for any increase in mass across the microfibril repeating unit using STEM mass mapping (Figure 5A). The microfibrils had a mean mass of ~3 MDa per repeat, which is slightly larger than the mass expected for 8 fibrillin molecules (~2.7 MDa). This suggests that some microfibril associated proteins (e.g. MAGP1 which is constitutively present) might have been co-purified. When LTBP-1 is present there is an increase in mass across the microfibril repeat of 350.2 ± 64.72 kDa which is localised predominantly to the bead region (Figure 5C). As the two major splice forms of LTBP-1 have molecular weights of 185 and 151 kDa respectively, this increase in mass is consistent with 2 molecules of LTBP-1 binding per microfibril repeat.

## Fibrillin arm region structure

To analyse further regions in the microfibril structure, the cryoEM dataset from the fibrillin microfibrils was recentred on the arm region and refined with a region-specific 3D mask. Particles which had been aligned to the full fibrillin repeat were recentred to locate the arm region coordinates at the particle centre. After 3D classification, the resulting best class was refined using 3D auto-refine with C2 symmetry imposed down the fibre axis, as with the bead reconstruction. The structure of the fibrillin arm region was determined, and it showed eight arms which are continuous from the bead to the interbead region and two shorter densities (Figure 6A). Domains from TB2 to TB3 were docked into this region guided by the domains located in the bead region and the antibody-mapping of mAb1919 (which recognises domains cbEGF7-Hybrid2). There are three distinct bands of higher density observed in the 2D averages of microfibrils which correlate with the docked locations of domains TB2, hybrid2 and TB3, which have a larger mass than the EGF/cbEGF domains (Figure 6C). Together, these data allow fitting of almost half of the fibrillin domains into the bead/arm region.

**Figure 6.**
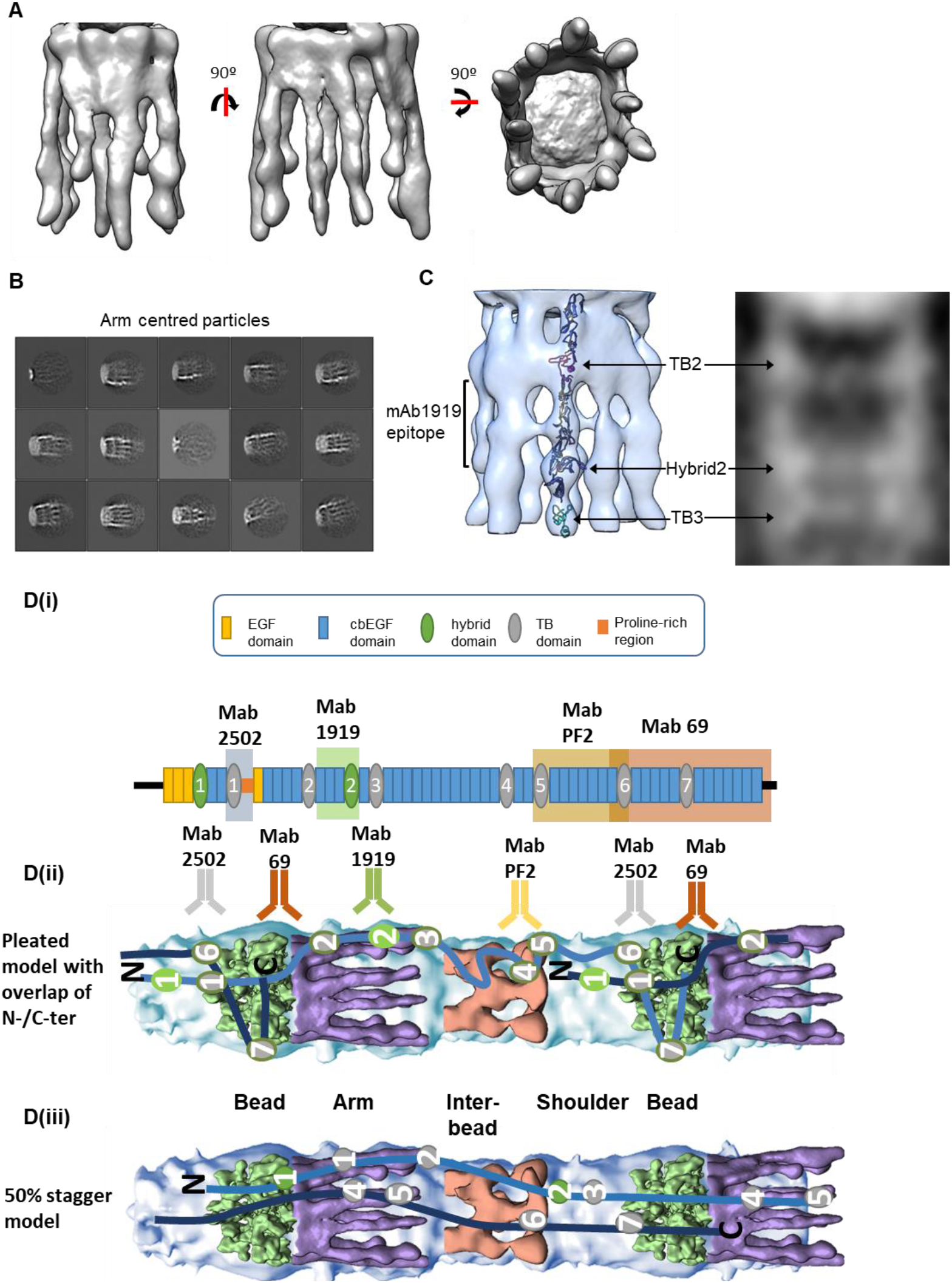
Fibrillin arm region structure. The cryoEM classes of the fibrillin microfibrils were recentred and refined to determine the structure of the fibrillin arm region. (A) CryoEM structure of the arm region of fibrillin microfibrils. (B) 2D class averages particles aligned to the fibrillin arm region reconstruction. Box size = 57 nm. (C) Domains docked into the arm region reconstruction from TB2 to TB3 where the larger TB and hybrid domains correspond with densities observed in EM images. (D) Schematic diagram of the intramolecular pleated and 50% staggered fibrillin molecular packing models. (Di) A domain schematic of fibrillin-1 which shows the epitopes for mabs 2502, 1919, 69 and PF2. A cartoon representation of the (ii) pleated model and (iii) 50% staggered model are shown superimposed over the negative stain structure of the full microfibril repeat (blue), and reconstructions of the interbead region (pink) [35] and the cryoEM structures of the bead (green) and arm (purple) regions. Overlapping fibrillin molecules are coloured light and dark blue alternately and the TB and hybrid domains are numbered. The epitopes for mabs 2502, 1919, 69 and PF2 are highlighted along the microfibril.

## Discussion

Here, we present the cryoEM structure of a native fibrillin microfibril, the first such analysis of a connective tissue-extracted fibrillar system. The microfibril was sub-divided into different regions for image analysis, to overcome flexibility along the microfibril axis, and the highest resolution was achieved for the bead region with sub-nm resolution. Guided by antibody-mapping data, we have docked an N-terminal region of the fibrillin molecule composed of domains TB1-cbEGF6 into the density which threads through the centre of the bead. Peptides from this region are rarely detected in MS analysis of microfibrils digested in non-denaturing conditions, which suggests that they are inaccessible to proteolytic enzymes and buried within the microfibril structure. When microfibrils are extracted in denaturing conditions, this region is present [42] which indicates that this region is present but buried and inaccessible in native conditions.

The C-terminal region, including TB7 and downstream cbEGF domains, could fit into the ring of density which wraps around the periphery of the bead (shown in grey in Figure 3E). This positioning would facilitate interleaving of the N- and C-terminal regions which would increases the density and mass of the bead region compared to interbead regions of the microfibril. This concentration of mass is consistent with STEM mass data and AFM height data for microfibrils [36]. An overlap of the N-and C-terminal regions is also supported by the positioning of a tissue transglutaminase cross-link between the N-terminal cbEGF5 domain and C-terminal TB7 domain [43]. The N-C-overlap predicted by this modal is also consistent with binding data from recombinant fibrillin fragments that show a C-terminal region (TB5-TB6) interacts with an N-terminal region [44].

The shoulder region was difficult to resolve with single particle averaging approaches suggesting that this region has conformational heterogeneity. Our model suggests that the fibrillin unique N-terminal or FUN domain resides in the shoulder region and NMR data have shown that this region is flexible and likely to be conformationally heterogeneous [45]. The shoulder region of the microfibril is also likely to be very compositionally heterogeneous as the N-terminal region binds to the majority of microfibril binding proteins (as reviewed in [2]).

Taken together, molecular modelling of the microfibril structure supports a pleated model, in which each fibrillin molecule is contained within each microfibril period with N- and C-terminal regions overlapping (Figure 6D). In this arrangement, the N-terminus of the fibrillin molecule is in the shoulder region, and the direction of the molecule runs through the bead region and into the arm and interbead regions. This molecular directionality continues through to the C-terminal domains which terminate at the next bead region and interact with the N-terminal region of an over-lapping fibrillin molecule. This is the axial path taken for one fibrillin molecule but there are eight molecules in cross-section forming each microfibril period [40]. This modelling is consistent with all published data on microfibrils including antibody-mapping [13, 36] and mass profile data [46]. Moreover, the position of the transglutaminase cross-link would be located in the bead region, perhaps giving rise to the inaccessible nature of the region containing cbEGF5 that forms the cross-link [43] (Supplementary Table). The pleated model fits better with the cryoEM density, microfibril mass features and antibody mapping data than the 50% stagger model (Figure 6D(ii)), but at this resolution we cannot rule out some intermediate packing arrangement.

To analyse the structural consequences of domain deletions on the microfibril and test the predictive capacity of the microfibril model, microfibrils from mice with deletions of the first hybrid domain and a WMS-causing deletion (lacking TB1-PRR-EGF4 domains) were analysed. Microfibrils containing either deletion had reduced periodicity, concomitant with the number of domains deleted, and showed disorganisation of the shoulder region. As the microfibril modelling would locate the first hybrid and TB1-PRR-EGF4 domains within or adjacent to the shoulder region, these data further support this microfibril modelling. The perturbation to the microfibril structure suggests that the mutant microfibrils may be less stable than the wildtype ones. Indeed, elastic fibres in the skin of WMS patients and mice carrying the WMS deletion have a moth-eaten appearance with abnormal aggregates of microfibrils [34].

The disruption of the microfibril structure was similar for both deletions, suggesting that both deletions may structurally perturb the conformation of neighbouring domains. Therefore, we hypothesised that the WMS deletion may disrupt binding to the first hybrid domain where latent TGFβ is known to bind via LTBP-1 [25]. Having confirmed that LTBP-1 binds the bead region of the microfibril, we also showed that the fibrillin-1 region containing the WMS deletion had reduced affinity for LTBP-1. However, TGFβ signalling does not appear to be disrupted in human WMS tissues or in WMS mouse models [34, 47] suggesting that the perturbation in binding does not directly contribute to the pathomechanism of disease.

Results from our study also significantly contribute to the current understanding of how structural alterations in fibrillin microfibril ultrastructure caused by fibrillin mutations leads to growth factor dysregulation. Currently, the bead region is seen as a critical microenvironment for growth factor sequestration and controlled activation [48, 49]. Any perturbation in this sensitive microenvironment leads to dysregulated growth and differentiation processes resulting in opposing clinical features (e.g. long bone overgrowth versus undergrowth, low muscle tone versus hypermuscularity) of the fibrillinopathies. We show that two LTBP-1 molecules can bind per bead unit. Given the length of LTBP-1 and the proximity of the Hybrid1 and FUN domains it is conceivable that binding of LTBP-1 may also restrict BMP binding or its activation. Furthermore, we show that the introduction of the WMS deletion mutation in the fibrillin molecule perturbs a firm LTBP-1 interaction to fibrillin. The structural rearrangement due to the WMS deletion mutation may prevent proper activation of sequestered BMPs from the microfibril scaffold as seen by us and others in WMS mouse models [47, 50].

When reconstituted *in vitro*, two molecules of LTBP-1 were able to bind per repeat to fibrillin microfibrils. Considering that there are eight fibrillin molecules per microfibril period [40], this suggests that the other sites are either already occupied or there is steric hindrance preventing more LTBP-1 molecules from binding. LTBP-1 is a large elongated molecule composed of more than 20 domains that can extend to ~40nm in length [51]. The mass increase upon LTBP-1-binding appears localised across and involves the centre of the bead, which suggests interactions can occur from the shoulder to the bead region, consistent with the elongated nature and length of LTBP-1. Therefore, there could be steric effects preventing further LTBP-1 molecules from binding. Furthermore, fibulins-2, −4 and −5 also bind to the first hybrid domain [25] which if co-purified with microfibrils could compete for the LTBP-1 binding site. Although MS data indicate that the fibulins are not major components in the ciliary zonule, and fibulin-2 is found in the vitreous [42]. However, a number of proteomics studies have shown that LTBP-2 is associated with microfibrils in ciliary zonules [38, 39, 42] which may compete with LTBP-1 for the available epitopes.

Indeed, in the reconstruction of the arm and bead regions, there are 10 arms that extend out from the bead. Eight of these are of similar size corresponding to the eight fibrillin molecules but two are shorter and terminate at the edge of the bead rather than continuing into the interbead (Supplementary figure 2). It is not clear whether the presence of these two truncated arms is an artefact of the 3D reconstruction due to flexibility of some of the arms or distortion of the structure by interaction with the air water interface, or whether the two smaller arms correspond to two microfibril binding proteins present in the microfibril structure. Indeed, STEM mass mapping performed on these microfibrils gave an experimental mass of ~3 MDa, which is larger than previously observed for microfibrils from canine ciliary zonules (2.55 MDa) [36]. Moreover, the expected mass of eight fibrillin molecules would be around 2.7 MDa suggesting microfibril-associated proteins may have co-purified in this preparation. MS data of ciliary zonule microfibrils after gel filtration show a number of LTBP-2 peptides (unpublished data) indicating that it co-purifies with these microfibrils, and LTBP-2 constitutes ~7% of proteins (fibrillin-1 ~75%) in the bovine zonule [42]. The LTBPs are members of the fibrillin superfamily and have a similar domain structure to fibrillin, so they would be expected to have a similar diameter and shape to fibrillin molecules, therefore, it is possible that microfibril-binding proteins have been observed in the 3D reconstruction of the microfibril.

Indeed, our purification does not use enzymatic digestion which may lead to a better preservation of the microfibril ultrastructure. It has been shown that microfibril isolation by enzymatic digestion destroys BMP binding epitopes [20]. Therefore, our new method allows us for the first time to investigate growth factor binding epitopes within the ultrastructure of the microfibril.

## Materials and methods

### Ethics statement

Isolation of fibrillin microfibrils from murine tissues was carried out in strict accordance with the German federal law on animal welfare, and the protocols were approved by the “Landesamt für Natur, Umwelt und Verbraucherschutz Nordrhein-Westfalen” for breeding (permit No. 84-02.04.2014.A397) and euthanasia (permit No. 84-02.05.40.14.115).

### Fibrillin microfibril extraction from bovine ciliary zonule

Ciliary zonules were extracted from dissected bovine ciliary zonule tissue and were disrupted using sonication in 20 mM Tris-HCl, 400 mM NaCl, 2 mM CaCl2, pH 7.4 with 2 x 10s pulses and cooled on ice for 30 seconds between pulses. Samples were centrifuged and the supernatant was separated using size exclusion chromatography with a 5ml CL-2B resin column. The column was equilibrated in either the sonication buffer or a physiological salt buffer for cryo-TEM studies (20 mM Tris-HCl, 150 mM NaCl 2 mM CaCl2 pH 7.4). The void volume containing fibrillin microfibrils was saved for further EM analysis.

### Fibrillin microfibril extraction from ΔH1 and WMS mice

Skin from 6 week old wildtype mice or homozygous for either deletion of the first hybrid domain (ΔH1) [33], or a deletion of three domains (TB1-PRR-EGF4) that causes Weill-Marchesani Syndrome (WMS)[34] was digested overnight at 4 °C in digestion buffer (0.1 mg/ml collagenase Type 1a (Sigma) in 20 mM Tris-HCl, 400 mM NaCl, 2 mM CaCl2 pH 7.4). Samples were centrifuged and separated using size exclusion chromatography as described above.

### Negative stain EM

Microfibrils extracted from mutant mouse skin were adsorbed onto glow discharged carbon coated copper grids (Agar) and were stained with 2% Uranyl acetate. Grids were imaged at a magnification of 23000x on a Tecnai G2 Polara TEM (FEI) equipped with a K2 Summit direct detector (Gatan) operating at an accelerating voltage of 300 kV. Images were collected over a 1 second exposure time in linear mode and were sampled at 1.4 Å/pixel. The microfibril repeat was manually picked in RELION [52] and subjected to 2D class averaging.

To determine the period length of the ΔH1 and WMS mutant microfibrils, the periodicity of 100 microfibril repeats were measured using ImageJ. Microfibrils were extracted from images and straightened using the straighten tool before measuring the repeat length.

### CryoEM sample preparation and data collection

Microfibril samples were adsorbed onto glow discharged holey carbon Quantifoil R1.2/1.3 grids before being blotted and plunge frozen in liquid ethane using a Vitrobot Mark IV (FEI). Microfibrils were imaged using automated data collection in EPU on a Titan Krios electron microscope (FEI) operating at an accelerating voltage of 300 kV. Movies comprising 20 frames with a 20 seconds exposure time and a total dose of 66 e Å^-2^ were collected on a K2 Summit direct detector (Gatan) in counting mode at a nominal magnification of 64000 which gave a pixel size of 2.2 Å. Initial imaging of microfibrils showed that they have a preferred orientation on the grid so during data collection the stage was tilted to 45 °.

### Single particle Data processing bead region reconstruction

Movies were motion corrected and dose weighted using MotionCorr2 (Zheng et al., 2017). Corrected images were imported into Warp [53] where 27,737 particles were picked using a Warp box net which had been previously trained on manually picked particles of the fibrillin microfibrils. Patch based CTF estimation was used to calculate local CTF values for the particles. Particle stacks were imported into cryoSPARC [54] and were used in a homogenous refinement using a negative stain reconstruction of the microfibril repeat as an initial model [35]. The resulting structure from cryoSPARC was then further refined in RELION 2.1 [52]. Particles which had been aligned to the full fibrillin repeat were recentred so the bead region or the arm region coordinates were then at the particle centre using a custom python script. For the bead region reconstruction, the shifted particle coordinates were then extracted and were 2D classified without alignment to remove bad particles; 13,184 particles from good classes were selected for 3D classification. The 3D classification of the shifted particles was performed with a restricted angular search using the command -- sigma_ang 5. The resulting best class was then further refined using 3D auto-refine with C2 symmetry imposed down the fibre axis. To remove any remaining bad particles a non-aligned 2D classification was performed and 7,139 particles were used in a final refinement. After post processing in RELION, the final structure had a resolution of 9.7 A.

For the arm region reconstruction, the shifted particle coordinates were also extracted and 2D classified without alignment to remove bad particles; 4957 good particles were selected for 3D classification. The 3D classification of the shifted particles was then performed with a restricted angular search using the command --sigma_ang 5. The resulting best class was refined using 3D auto-refine with C2 symmetry imposed down the fibre axis as with the bead reconstruction. After post processing in RELION the final arm structure had a resolution of 18.3 Å (Supplementary figure 3). The EM data has been deposited to EMBD with accession codes EMD-13984 for the bead model and EMD-13986 for the arm region.

### Refining antibody epitopes with recombinant fibrillin fragments

Recombinant fibrillin fragments were expressed and purified as previously described [19, 55]. Proteins were detected by western blotting with antibodies mab2502 (clone 26) and mab1919 (clone 11C1.3) from Sigma.

### LTBP-1 binding and STEM mass mapping

Full-length LTBP-1 was purified as previously described [51] and incubated with fibrillin microfibrils in a 2:1 molar ratio for 4 hours at 4 °C. The complex was adhered for 60 seconds to 400-mesh carbon coated grids then washed with milliQ H2O three times and dried. The sample was visualised in STEM mode with a Fishione high-angle annular dark-field detector on a FEI Tecnai12 Twin TEM at 34,000x magnification. A camera length of 350 cm was used to give an angular collection range of 20–100 mrad. Tobacco mosaic virus was used as a calibration standard for mass per unit length. The electron dose was kept sufficiently low (< 300 e.nm^-1^) to produce negligible mass loss. Mass per unit length measurements and axial mass distributions were measured from STEM ADF images using the Semper6 image analysis software (Synoptics, Cambridge, UK).

### Fibrillin-LTBP-1 binding by surface plasmon resonance

The C-terminal region of LTBP-1 was purified as previously described [51] and immobilised by amine coupling onto a CM5 sensor chip at 2.5 μg/ml in 50 mM sodium acetate buffer, pH 3.0 in a BIAcore T200 biosensor; typically 200 response units (RU) were immobilised. Recombinant fibrillin-1 fragments PF3 and PF3 WMS (0-50 nM) were injected onto the sensor chip at a flow rate of 50 μl/min in 10 mM HEPES, 150 mM NaCl, 1 mM CalCl2, 0.05% Tween-20 for 100 seconds and then allowed to dissociate for 180 seconds. Regeneration was performed by injection of 10 mM glycine, pH 2 for 30 seconds at a flow rate of 30 μL/min followed by a stabilisation period of 60 seconds. KDs were calculated using equilibrium analysis: equilibrium response was plotted against concentration and non-linear regression was used to calculate KD using the equation for one-site binding. Each assay was performed at least twice.

## Acknowledgements

We would like to thank staff in the EM Facility (RRID:SCR_021147; Faculty of Biology, Medicine and Health, University of Manchester) for assistance, and funding from the BBSRC (ref: BB/T017643/1) for the Glacios cryoEM used for screening. The Wellcome Centre for Cell-Matrix Research is supported by funding from the Wellcome Trust (203128/Z/16/Z). A.R.F.G. is supported by BBSRC funding (Ref: BB/N015398/1 and BB/S015779/1). We acknowledge the UK national electron bio-imaging centre (eBIC) for access to cryoEM facilities (ref: EM16619-5). Data were collected at the Astbury Biostructure Laboratory on the FEI Titan Krios microscopes which were funded by the University of Leeds and Wellcome Trust (108466/Z/15/Z). Funding for this study was also provided by the Deutsche Forschungsgemeinschaft (DFG, German Research Foundation) project numbers 73111208 (SFB 829/B12), 384170921 FOR2722/ C2, and 397484323 TRR259/ B09 to G.S.

## References

[1] Godwin ARF, Singh M, Lockhart-Cairns MP, Alanazi YF, Cain SA, Baldock C. The role of fibrillin and microfibril binding proteins in elastin and elastic fibre assembly. Matrix Biol. 2019;84:17–30.

[2] Thomson J, Singh M, Eckersley A, Cain SA, Sherratt MJ, Baldock C. Fibrillin microfibrils and elastic fibre proteins: Functional interactions and extracellular regulation of growth factors. Semin Cell Dev Biol. 2019;89:109–17.

[3] Pereira L, D’Alessio M, Ramirez F, Lynch JR, Sykes B, Pangilinan T, et al. Genomic organization of the sequence coding for fibrillin, the defective gene product in Marfan syndrome. Hum Mol Genet. 1993;2:961–8.

[4] Zhang H, Apfelroth SD, Hu W, Davis EC, Sanguineti C, Bonadio J, et al. Structure and expression of fibrillin-2, a novel microfibrillar component preferentially located in elastic matrices. J Cell Biol. 1994;124:855–63.

[5] Corson GM, Charbonneau NL, Keene DR, Sakai LY. Differential expression of fibrillin-3 adds to microfibril variety in human and avian, but not rodent, connective tissues. Genomics. 2004;83:461–72.

[6] Maslen CL, Corson GM, Maddox BK, Glanville RW, Sakai LY. Partial sequence of a candidate gene for the Marfan syndrome. Nature. 1991;352:334–7.

[7] Kanzaki T, Olofsson A, Moren A, Wernstedt C, Hellman U, Miyazono K, et al. TGF-beta 1 binding protein: a component of the large latent complex of TGF-beta 1 with multiple repeat sequences. Cell. 1990;61:1051–61.

[8] Moren A, Olofsson A, Stenman G, Sahlin P, Kanzaki T, Claesson-Welsh L, et al. Identification and characterization of LTBP-2, a novel latent transforming growth factor-beta-binding protein. J Biol Chem. 1994;269:32469–78.

[9] Yin W, Smiley E, Germiller J, Mecham RP, Florer JB, Wenstrup RJ, et al. Isolation of a novel latent transforming growth factor-beta binding protein gene (LTBP-3). J Biol Chem. 1995;270:10147–60.

[10] Giltay R, Kostka G, Timpl R. Sequence and expression of a novel member (LTBP-4) of the family of latent transforming growth factor-beta binding proteins. FEBS Lett. 1997;411:164–8.

[11] Keene DR, Maddox BK, Kuo HJ, Sakai LY, Glanville RW. Extraction of extendable beaded structures and their identification as fibrillin-containing extracellular matrix microfibrils. The journal of histochemistry and cytochemistry: official journal of the Histochemistry Society. 1991;39:441–9.

[12] Kielty CM, Sherratt MJ, Marson A, Baldock C. Fibrillin microfibrils. Advances in protein chemistry. 2005;70:405–36.

[13] Reinhardt DP, Keene DR, Corson GM, Poschl E, Bachinger HP, Gambee JE, et al. Fibrillin-1: organization in microfibrils and structural properties. J Mol Biol. 1996;258:104–16.

[14] Lin G, Tiedemann K, Vollbrandt T, Peters H, Batge B, Brinckmann J, et al. Homo-and heterotypic fibrillin-1 and −2 interactions constitute the basis for the assembly of microfibrils. J Biol Chem. 2002;277:50795–804.

[15] Marson A, Rock MJ, Cain SA, Freeman LJ, Morgan A, Mellody K, et al. Homotypic fibrillin-1 interactions in microfibril assembly. J Biol Chem. 2005;280:5013–21.

[16] Hubmacher D, El-Hallous EI, Nelea V, Kaartinen MT, Lee ER, Reinhardt DP. Biogenesis of extracellular microfibrils: Multimerization of the fibrillin-1 C terminus into bead-like structures enables self-assembly. Proceedings of the National Academy of Sciences of the United States of America. 2008;105:6548–53.

[17] Sherratt MJ, Wess TJ, Baldock C, Ashworth J, Purslow PP, Shuttleworth CA, et al. Fibrillin-rich microfibrils of the extracellular matrix: ultrastructure and assembly. Micron (Oxford, England: 1993). 2001;32:185–200.

[18] Lu Y, Sherratt MJ, Wang MC, Baldock C. Tissue specific differences in fibrillin microfibrils analysed using single particle image analysis. Journal of structural biology. 2006;155:285–93.

[19] Baldock C, Siegler V, Bax DV, Cain SA, Mellody KT, Marson A, et al. Nanostructure of fibrillin-1 reveals compact conformation of EGF arrays and mechanism for extensibility. Proceedings of the National Academy of Sciences of the United States of America. 2006;103:11922–7.

[20] Kuo CL, Isogai Z, Keene DR, Hazeki N, Ono RN, Sengle G, et al. Effects of fibrillin-1 degradation on microfibril ultrastructure. J Biol Chem. 2007;282:4007–20.

[21] Faivre L, Gorlin RJ, Wirtz MK, Godfrey M, Dagoneau N, Samples JR, et al. In frame fibrillin-1 gene deletion in autosomal dominant Weill-Marchesani syndrome. Journal of medical genetics. 2003;40:34–6.

[22] Bax DV, Bernard SE, Lomas A, Morgan A, Humphries J, Shuttleworth CA, et al. Cell adhesion to fibrillin-1 molecules and microfibrils is mediated by alpha 5 beta 1 and alpha v beta 3 integrins. J Biol Chem. 2003;278:34605–16.

[23] Jovanovic J, Takagi J, Choulier L, Abrescia NG, Stuart DI, van der Merwe PA, et al. alphaVbeta6 is a novel receptor for human fibrillin-1. Comparative studies of molecular determinants underlying integrin-rgd affinity and specificity. J Biol Chem. 2007;282:6743–51.

[24] Bax DV, Mahalingam Y, Cain S, Mellody K, Freeman L, Younger K, et al. Cell adhesion to fibrillin-1: identification of an Arg-Gly-Asp-dependent synergy region and a heparin-binding site that regulates focal adhesion formation. J Cell Sci. 2007;120:1383–92.

[25] Ono RN, Sengle G, Charbonneau NL, Carlberg V, Bachinger HP, Sasaki T, et al. Latent transforming growth factor beta-binding proteins and fibulins compete for fibrillin-1 and exhibit exquisite specificities in binding sites. J Biol Chem. 2009;284:16872–81.

[26] Sengle G, Charbonneau NL, Ono RN, Sasaki T, Alvarez J, Keene DR, et al. Targeting of bone morphogenetic protein growth factor complexes to fibrillin. J Biol Chem. 2008;283:13874–88.

[27] Sengle G, Ono RN, Sasaki T, Sakai LY. Prodomains of transforming growth factor beta (TGFbeta) superfamily members specify different functions: extracellular matrix interactions and growth factor bioavailability. J Biol Chem. 2011;286:5087–99.

[28] Miyazono K, Olofsson A, Colosetti P, Heldin CH. A role of the latent TGF-beta 1-binding protein in the assembly and secretion of TGF-beta 1. EMBO J. 1991;10:1091–101.

[29] Gleizes PE, Beavis RC, Mazzieri R, Shen B, Rifkin DB. Identification and characterization of an eight-cysteine repeat of the latent transforming growth factor-beta binding protein-1 that mediates bonding to the latent transforming growth factor-beta1. J Biol Chem. 1996;271:29891–6.

[30] Saharinen J, Taipale J, Keski-Oja J. Association of the small latent transforming growth factor-beta with an eight cysteine repeat of its binding protein LTBP-1. The EMBO journal. 1996;15:245–53.

[31] Chen Y, Ali T, Todorovic V, O’Leary J M, Kristina Downing A, Rifkin DB. Amino acid requirements for formation of the TGF-beta-latent TGF-beta binding protein complexes. J Mol Biol. 2005;345:175–86.

[32] Robinson PN, Arteaga-Solis E, Baldock C, Collod-Beroud G, Booms P, De Paepe A, et al. The molecular genetics of Marfan syndrome and related disorders. Journal of medical genetics. 2006;43:769–87.

[33] Charbonneau NL, Carlson EJ, Tufa S, Sengle G, Manalo EC, Carlberg VM, et al. In vivo studies of mutant fibrillin-1 microfibrils. J Biol Chem. 2010;285:24943–55.

[34] Sengle G, Tsutsui K, Keene DR, Tufa SF, Carlson EJ, Charbonneau NL, et al. Microenvironmental regulation by fibrillin-1. PLoS genetics. 2012;8:e1002425.

[35] Godwin ARF, Starborg T, Smith DJ, Sherratt MJ, Roseman AM, Baldock C. Multiscale Imaging Reveals the Hierarchical Organization of Fibrillin Microfibrils. J Mol Biol. 2018;430:4142–55.

[36] Baldock C, Koster AJ, Ziese U, Rock MJ, Sherratt MJ, Kadler KE, et al. The supramolecular organization of fibrillin-rich microfibrils. J Cell Biol. 2001;152:1045–56.

[37] Maddox BK, Sakai LY, Keene DR, Glanville RW. Connective tissue microfibrils. Isolation and characterization of three large pepsin-resistant domains of fibrillin. J Biol Chem. 1989;264:21381–5.

[38] Eckersley A, Mellody KT, Pilkington S, Griffiths CEM, Watson REB, O’Cualain R, et al. Structural and compositional diversity of fibrillin microfibrils in human tissues. J Biol Chem. 2018;293:5117–33.

[39] Cain SA, Morgan A, Sherratt MJ, Ball SG, Shuttleworth CA, Kielty CM. Proteomic analysis of fibrillin-rich microfibrils. Proteomics. 2006;6:111–22.

[40] Wang MC, Lu Y, Baldock C. Fibrillin microfibrils: a key role for the interbead region in elasticity. J Mol Biol. 2009;388:168–79.

[41] Lockhart-Cairns MP, Newandee H, Thomson J, Weiss AS, Baldock C, Tarakanova A. Transglutaminase-Mediated Cross-Linking of Tropoelastin to Fibrillin Stabilises the Elastin Precursor Prior to Elastic Fibre Assembly. Journal of Molecular Biology. 2020;432:5736–51.

[42] De Maria A, Wilmarth PA, David LL, Bassnett S. Proteomic Analysis of the Bovine and Human Ciliary Zonule. Investigative ophthalmology & visual science. 2017;58:573–85.

[43] Qian RQ, Glanville RW. Alignment of fibrillin molecules in elastic microfibrils is defined by transglutaminase-derived cross-links. Biochemistry. 1997;36:15841–7.

[44] Chaudhry SS, Cain SA, Morgan A, Dallas SL, Shuttleworth CA, Kielty CM. Fibrillin-1 regulates the bioavailability of TGFbeta1. J Cell Biol. 2007;176:355–67.

[45] Yadin DA, Robertson IB, McNaught-Davis J, Evans P, Stoddart D, Handford PA, et al. Structure of the fibrillin-1 N-terminal domains suggests that heparan sulfate regulates the early stages of microfibril assembly. Structure. 2013;21:1743–56.

[46] Sherratt MJ, Holmes DF, Shuttleworth CA, Kielty CM. Scanning transmission electron microscopy mass analysis of fibrillin-containing microfibrils from foetal elastic tissues. The international journal of biochemistry & cell biology. 1997;29:1063–70.

[47] Mularczyk EJ, Singh M, Godwin ARF, Galli F, Humphreys N, Adamson AD, et al. ADAMTS10-mediated tissue disruption in Weill-Marchesani syndrome. Hum Mol Genet. 2018;27:3675–87.

[48] Sengle G, Sakai LY. The fibrillin microfibril scaffold: A niche for growth factors and mechanosensation? Matrix Biol. 2015;47:3–12.

[49] Zigrino P, Sengle G. Fibrillin microfibrils and proteases, key integrators of fibrotic pathways. Adv Drug Deliv Rev. 2019;146:3–16.

[50] Oichi T, Taniguchi Y, Soma K, Oshima Y, Yano F, Mori Y, et al. Adamts17 is involved in skeletogenesis through modulation of BMP-Smad1/5/8 pathway. Cell Mol Life Sci. 2019;76:4795–809.

[51] Troilo H, Steer R, Collins RF, Kielty CM, Baldock C. Independent multimerization of Latent TGFbeta Binding Protein-1 stabilized by cross-linking and enhanced by heparan sulfate. Sci Rep. 2016;6:34347.

[52] Kimanius D, Forsberg BO, Scheres SHW, Lindahl E. Accelerated cryo-EM structure determination with parallelisation using GPUs in RELION-2. eLife. 2016;5:e18722.

[53] Tegunov D, Cramer P. Real-time cryo-electron microscopy data preprocessing with Warp. Nature methods. 2019;16:1146–52.

[54] Punjani A, Rubinstein JL, Fleet DJ, Brubaker MA. cryoSPARC: algorithms for rapid unsupervised cryo-EM structure determination. Nature methods. 2017;14:290–6.

[55] Cain SA, Baldwin AK, Mahalingam Y, Raynal B, Jowitt TA, Shuttleworth CA, et al. Heparan sulfate regulates fibrillin-1 N-and C-terminal interactions. J Biol Chem. 2008;283:27017–27.

