## Supplementary figures for "Fibrillin microfibril structure identifies long-range effects of inherited pathogenic mutations affecting a key regulatory TGFβ-binding site"

### Slide 1
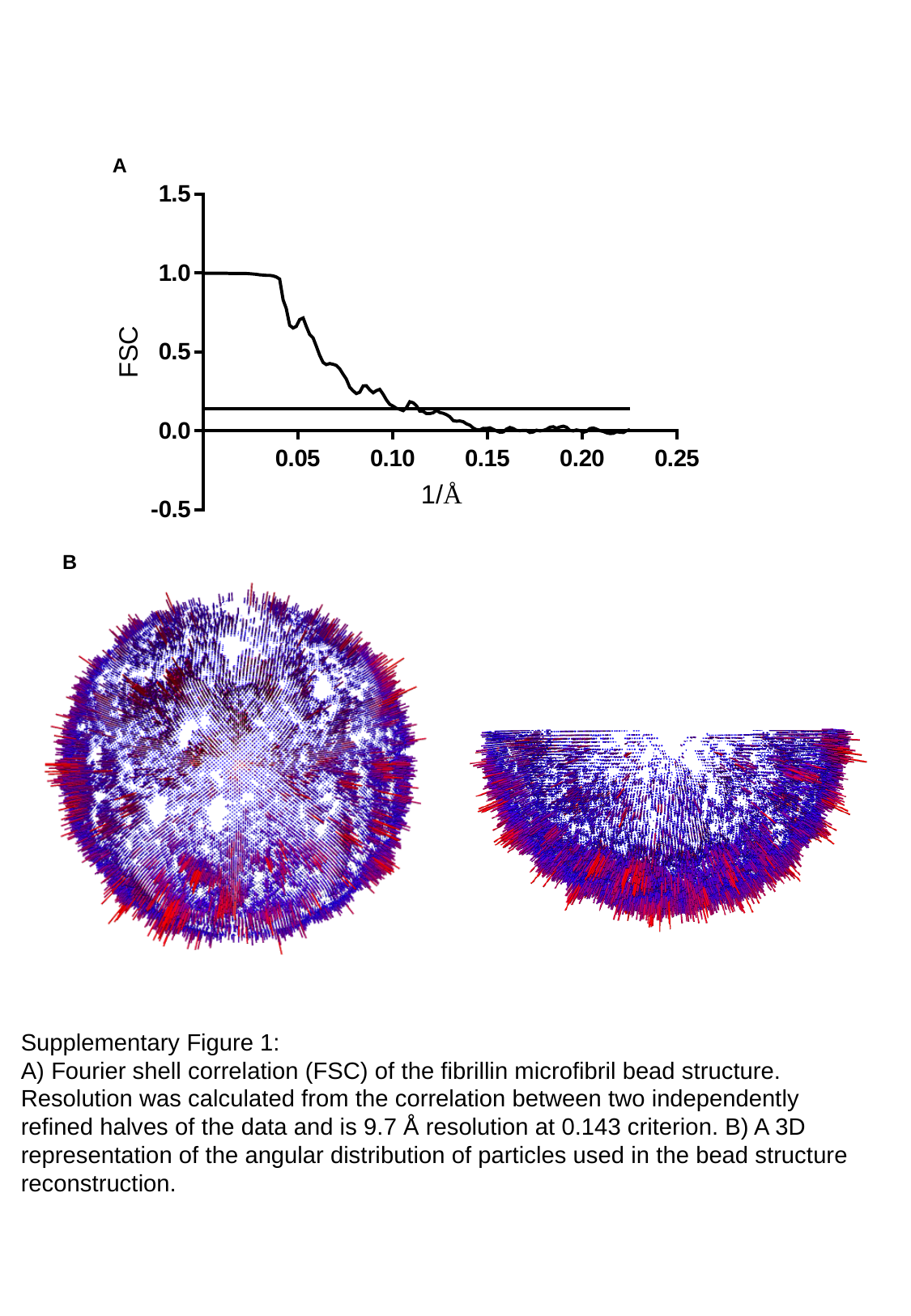

A
B
Supplementary Figure 1:
A) Fourier shell correlation (FSC) of the fibrillin microfibril bead structure. Resolution was calculated from the correlation between two independently refined halves of the data and is 9.7 Å resolution at 0.143 criterion. B) A 3D representation of the angular distribution of particles used in the bead structure reconstruction.

### Slide 2
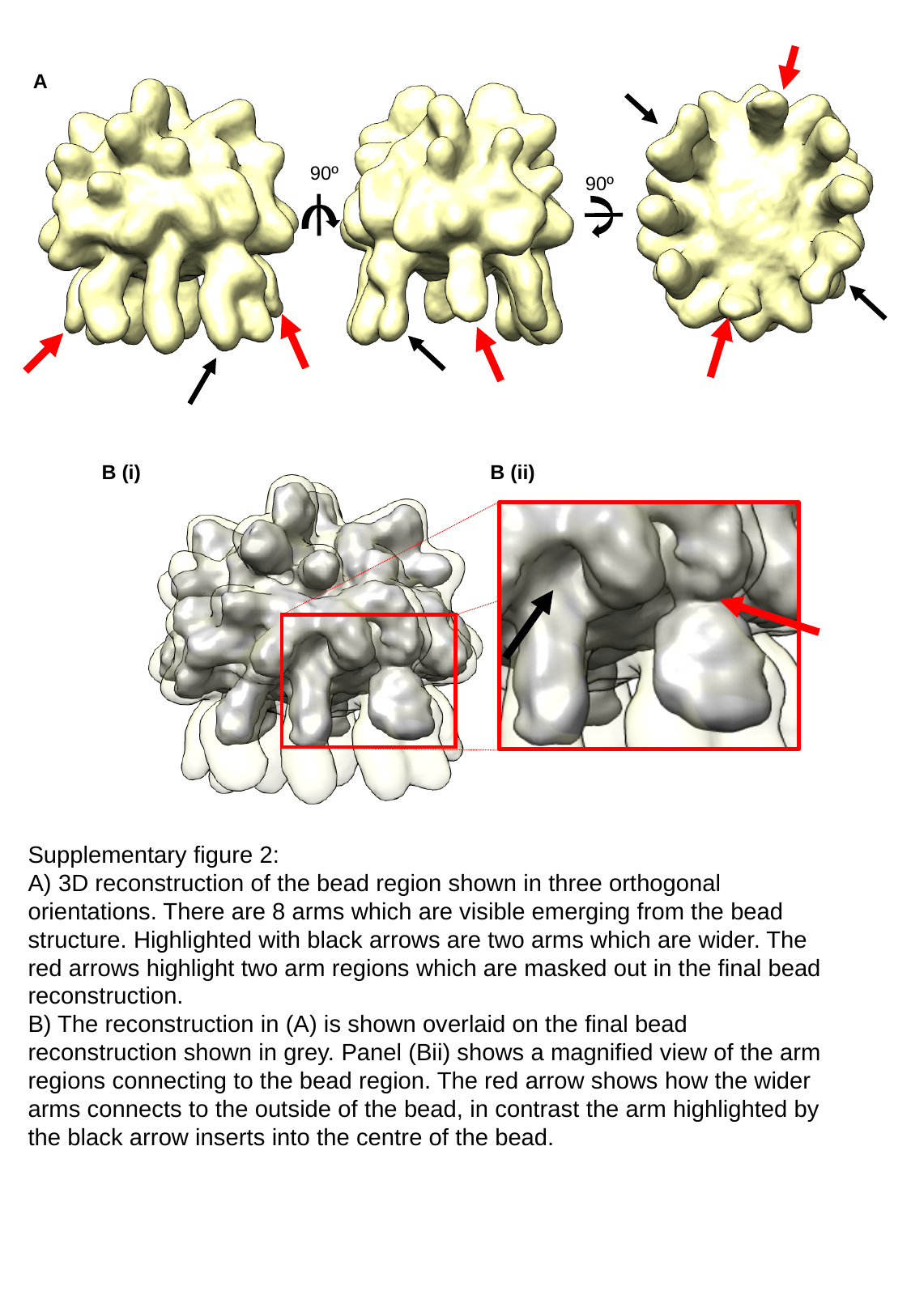

A
90º
90º
B (i)
B (ii)
Supplementary figure 2:
A) 3D reconstruction of the bead region shown in three orthogonal orientations. There are 8 arms which are visible emerging from the bead structure. Highlighted with black arrows are two arms which are wider. The red arrows highlight two arm regions which are masked out in the final bead reconstruction.
B) The reconstruction in (A) is shown overlaid on the final bead reconstruction shown in grey. Panel (Bii) shows a magnified view of the arm regions connecting to the bead region. The red arrow shows how the wider arms connects to the outside of the bead, in contrast the arm highlighted by the black arrow inserts into the centre of the bead.

### Slide 3
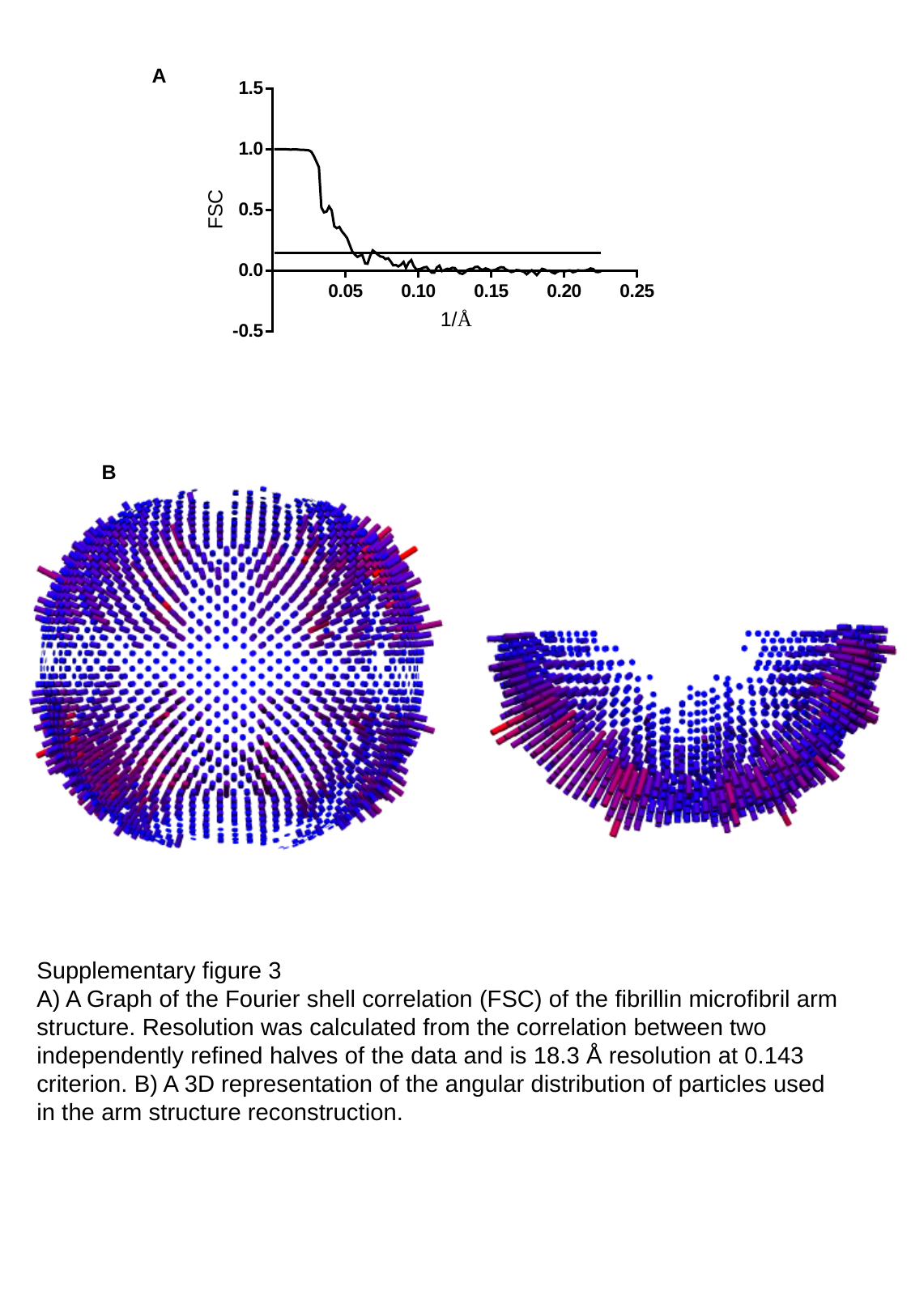

A
B
Supplementary figure 3
A) A Graph of the Fourier shell correlation (FSC) of the fibrillin microfibril arm structure. Resolution was calculated from the correlation between two independently refined halves of the data and is 18.3 Å resolution at 0.143 criterion. B) A 3D representation of the angular distribution of particles used in the arm structure reconstruction.

### Slide 4
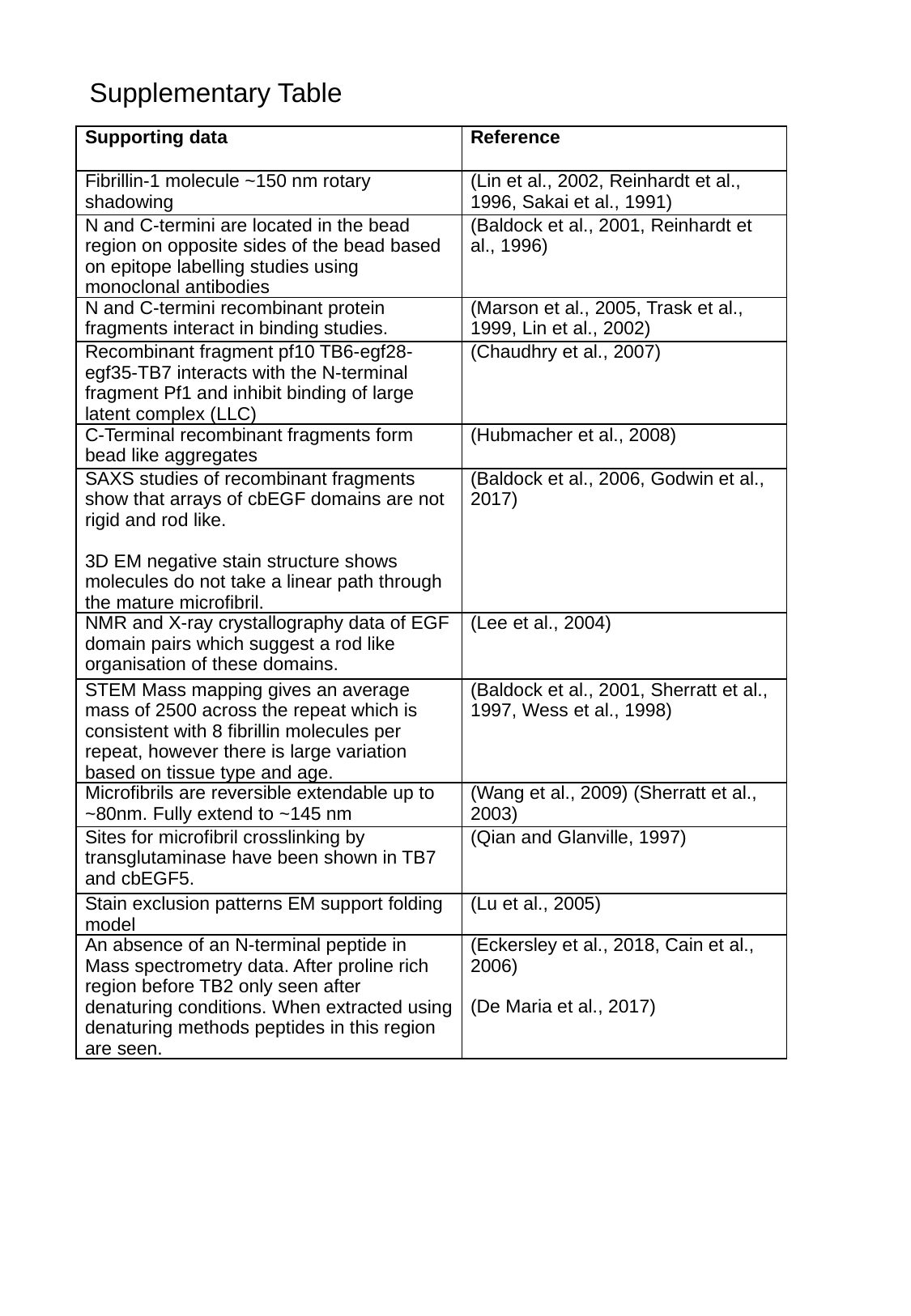

Supplementary Table
| Supporting data | Reference |
| --- | --- |
| Fibrillin-1 molecule ~150 nm rotary shadowing | (Lin et al., 2002, Reinhardt et al., 1996, Sakai et al., 1991) |
| N and C-termini are located in the bead region on opposite sides of the bead based on epitope labelling studies using monoclonal antibodies | (Baldock et al., 2001, Reinhardt et al., 1996) |
| N and C-termini recombinant protein fragments interact in binding studies. | (Marson et al., 2005, Trask et al., 1999, Lin et al., 2002) |
| Recombinant fragment pf10 TB6-egf28-egf35-TB7 interacts with the N-terminal fragment Pf1 and inhibit binding of large latent complex (LLC) | (Chaudhry et al., 2007) |
| C-Terminal recombinant fragments form bead like aggregates | (Hubmacher et al., 2008) |
| SAXS studies of recombinant fragments show that arrays of cbEGF domains are not rigid and rod like. 3D EM negative stain structure shows molecules do not take a linear path through the mature microfibril. | (Baldock et al., 2006, Godwin et al., 2017) |
| NMR and X-ray crystallography data of EGF domain pairs which suggest a rod like organisation of these domains. | (Lee et al., 2004) |
| STEM Mass mapping gives an average mass of 2500 across the repeat which is consistent with 8 fibrillin molecules per repeat, however there is large variation based on tissue type and age. | (Baldock et al., 2001, Sherratt et al., 1997, Wess et al., 1998) |
| Microfibrils are reversible extendable up to ~80nm. Fully extend to ~145 nm | (Wang et al., 2009) (Sherratt et al., 2003) |
| Sites for microfibril crosslinking by transglutaminase have been shown in TB7 and cbEGF5. | (Qian and Glanville, 1997) |
| Stain exclusion patterns EM support folding model | (Lu et al., 2005) |
| An absence of an N-terminal peptide in Mass spectrometry data. After proline rich region before TB2 only seen after denaturing conditions. When extracted using denaturing methods peptides in this region are seen. | (Eckersley et al., 2018, Cain et al., 2006) (De Maria et al., 2017) |
